# Neurodevelopmental disorder associated mutations in TAOK1 reveal its function as a plasma membrane remodeling kinase

**DOI:** 10.1101/2022.07.06.499025

**Authors:** Neal Beeman, Tanmay Sapre, Shao-En Ong, Smita Yadav

## Abstract

Mutations in *TAOK1*, a serine-threonine kinase encoding gene are strongly associated with both autism spectrum disorder (ASD) and neurodevelopmental delay (NDD). However, molecular function of this evolutionarily conserved kinase and the mechanisms through which *TAOK1* mutations lead to neuropathology are unknown. Here, we showed that TAOK1 is highly expressed in neurons within the brain, and has a functional role in remodeling the plasma membrane through direct association with phosphoinositides. We characterized four NDD-associated TAOK1 mutations, and demonstrated that these mutations render TAOK1 catalytically dead. Kinase dead TAOK1 mutants were aberrantly trapped in membrane-bound state, which induced exuberant membrane protrusions. Expression of TAOK1 disease mutants in hippocampal neurons led to abnormal growth of the dendritic arbor. The coiled-coil region C-terminal to the kinase domain are predicted to fold into a triple helix. We showed that this triple helix directly bound phospholipids, and was required for both TAOK1 membrane association and induction of aberrant protrusions. Further, TAOK1 mutants were rescued from their membrane-trapped state by exogenous expression of the isolated kinase domain. Utilizing mass-spectrometry, we identified critical residues in the triple helix phosphorylated by TAOK1 that autoregulated its plasma membrane association. These findings define a previously unknown function of TAOK1 as a unique plasma membrane remodeling kinase, and reveal the underlying mechanisms through which TAOK1 dysfunction leads to neurodevelopmental disorders.

## INTRODUCTION

Neuronal morphogenesis, migration, and maturation rely on extensive remodeling of the neuronal plasma membrane. In addition to the well-appreciated role of the microtubule and actin cytoskeleton in regulating plasma membrane shape and dynamics, proteins can directly associate with the cell membrane through electrostatic interactions to deform and shape membrane (Doherty and McMahon, 2008). Membrane remodeling proteins identified to date belong to the Bin/Amphiphysin/Rvs (BAR) superfamily which consist of a membrane-associating BAR domain. The BAR domain is structurally defined as an elongated homodimer with a shallow curved surface formed by the antiparallel interaction of two triple helix coiled-coils (Frost et al., 2009, 2008; Heath and Insall, 2008; Peter et al., 2004). This curved BAR dimer interacts with cellular membranes and causes membrane deformation (Henne et al., 2007). Depending on whether the concave or convex surface of the BAR dimer binds membranes, either membrane invagination or outward protrusions are generated (Heath and Insall, 2008; Mattila et al., 2007; Saarikangas et al., 2009). To date, membrane remodeling proteins outside the BAR domain family have not been identified. In this study, we report that the autism risk gene, *TAOK1*, encodes a unique protein kinase that can remodel the plasma membrane by direct binding with phospholipids in an activity dependent manner.

Thousand and one amino acid kinase 1 (TAOK1) is an evolutionarily conserved serine-threonine kinase (Moore et al., 2000; Zihni et al., 2006). Recently, several missense and truncation mutations in *TAOK1* have been recently identified in patients with severe neurodevelopmental delay (NDD) (Dulovic-Mahlow et al., 2019; Hunter et al., 2022; Satterstrom et al., 2020; Woerden et al., 2021). The core clinical features of patients with TAOK1-associated NDD syndrome include varying degrees of intellectual disability and developmental delay, neonatal feeding difficulties, overlapping dysmorphic facial features, behavior problems, hypotonia, and joint hypermobility (Dulovic-Mahlow et al., 2019; Hunter et al., 2022; Woerden et al., 2021). While evidence for the disease association of TAOK1 is compelling, little is known about the molecular and cellular function of TAOK1 in neuronal development. Knockdown of TAOK1 *in utero* in mice causes defects in neuronal migration, and its overexpression leads to dendritic arborization defects (Dulovic-Mahlow et al., 2019). TAOK1 is phosphorylated by hippo kinase MST3, which is important for dendritic spine formation (Ultanir et al., 2014). In *D. melanogaster*, activity of homolog dTao is required for neuroblast proliferation and neuronal connectivity (Hu et al., 2020; Poon et al., 2016; Tenedini et al., 2019). Kin-18 is a homolog of Tao kinases in *C. elegans* that is important for development of embryonic polarity (Spiga et al., 2013). Despite its role in diverse aspects of neuronal development and its strong disease association, the molecular function of TAOK1 is unknown. Further, how mutations associated with NDD impact TAOK1 function and contribute to neuropathology remains unknown.

Through the study of autism-associated mutations in TAOK1, we uncovered a hitherto unknown function of TAOK1 as a plasma membrane remodeling kinase. We demonstrate its remarkable property to bind phosphoinositides and induce plasma membrane protrusions in an activity dependent manner. Kinase-dead NDD-associated mutations disrupt the autoregulation of TAOK1 resulting in constitutive membrane binding that leads to aberrant neuronal membrane protrusions and dendritic growth defects in hippocampal neurons. By delineating how TAOK1 controls its membrane association, our results provide evidence for a previously unappreciated regulatory mechanism for membrane remodeling that is important for neuronal morphogenesis, the dysfunction of which contributes to neuropathology in autism and neurodevelopmental delay.

## RESULTS

### *TAOK1* is highly expressed in neurons in the mammalian brain

TAOK1 is a large protein of 1001 amino acids that comprises an N-terminal kinase domain followed by predicted coiled-coil domains of unknown function. Both the kinase domain and the coiled-coils are structurally conserved across worm, fly, fish, rodent and human species (Figure 1A). We first investigated expression level of *TAOK1* mRNA in the brain in both human and rodents by analyzing transcriptome databases (Kang et al., 2011; Sjöstedt et al., 2020; Sunkin et al., 2013). In humans, *TAOK1* is highly expressed in the prenatal and postnatal stages of brain development, and in the adult human across brain regions including neocortex, hippocampus, cerebellum, amygdala and striatum (Figure S1A-E). At the protein level, TAOK1 was the highest in neuronal cell types in both human cortex and hippocampus, with mild or low protein detected in non-neuronal cells (Figure S1F). Similarly, TAOK1 mRNA was widely expressed in all brain regions in the mouse brain. Cortical layer 2/3, the pyramidal layer hippocampal CA1, granular layer CA3 and DG and cerebellum exhibited the highest expression by *in situ* hybridization (Figure 1B). Using antibodies against TAOK1 protein, we found that TAOK1 was expressed at a higher level in hippocampal neurons compared to non-neuronal cells and was localized throughout the soma, dendrites and axons (Figure 1C-D). Next, we generated a GFP-tagged TAOK1 construct, which upon expression in hippocampal neurons was localized similarly to the endogenous TAOK1 protein and was present in all neuronal subcompartments (Figure 1D).

**Figure 1.**
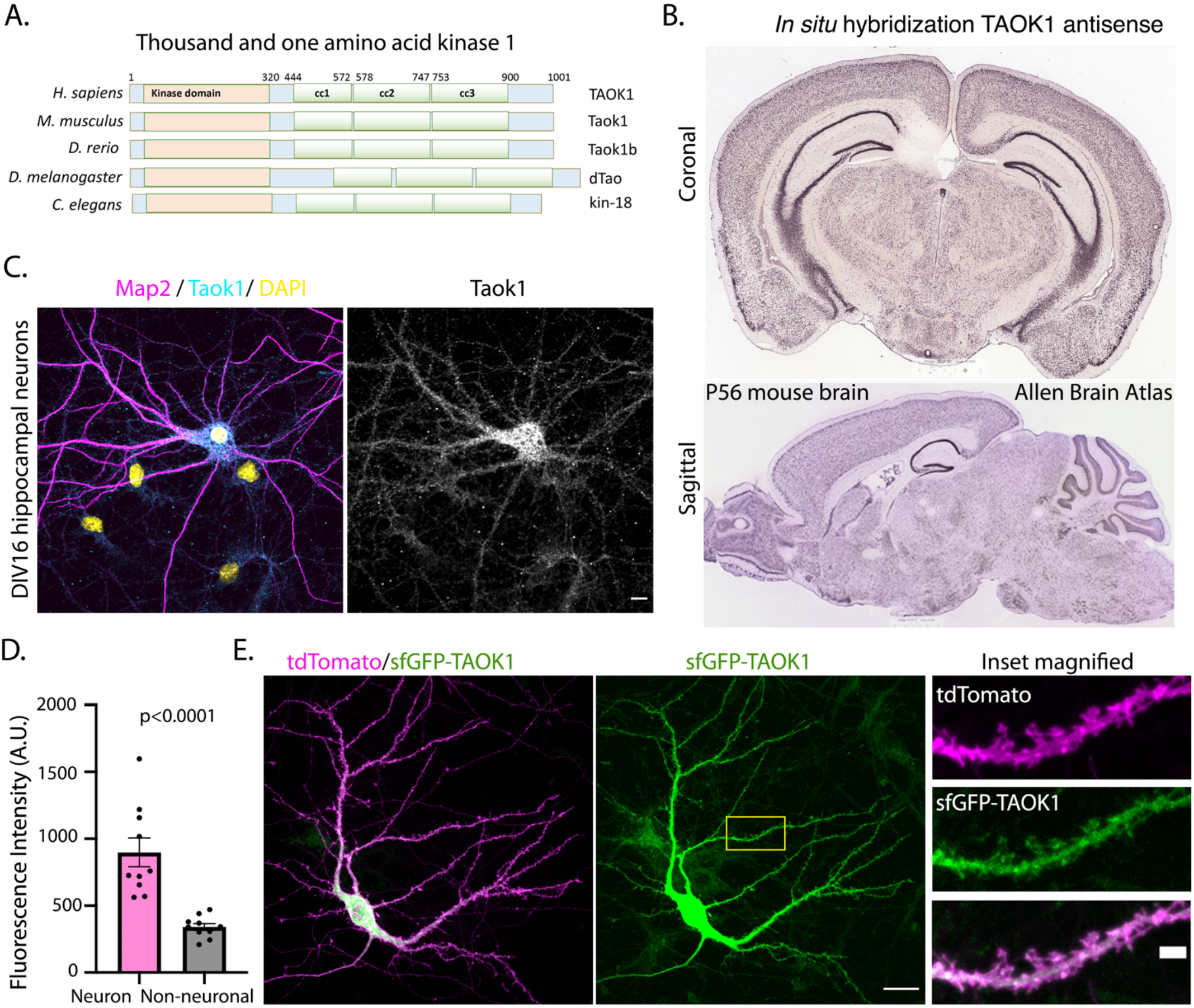
TAOK1 is an evolutionarily conserved serine-threonine kinase that is highly expressed in hippocampal neurons. (A) Schematic representation of secondary structure of Thousand and one amino acid kinase 1 orthologs found in human (TAOK1), mouse (Taok1), fish (Taok1b), fly (dTao) and worm (kin-18) species. The N-terminal kinase domain (orange) and three predicted coiled-coils (green) are highly evolutionarily conserved at the sequence and structural level. (B) *In situ* hybridization data from Allen Brain Atlas using antisense probe against Taok1 on sagittal and coronal sections of P56 C57BL6/J male mouse brain. (C) Primary hippocampal neurons obtained from E15 rat embryos were fixed at 16 days *in vitro* (DIV) and immunostained using antibodies against Taok1 (cyan) and dendritic protein Map2 (magenta). Nucleus is stained with DAPI (yellow). Grayscale image shows Taok1 immunostaining. Scale bar is 10µm. (D) Immunofluorescence intensity of Taok1 in neuronal cells compared to non-neuronal cells (n= 10 neuronal (Map2 positive) and non-neuronal cells (Map2 negative), p<0.0001). (E) DIV9 hippocampal neurons were transfected with sfGFP-TAOK1 (green) along with cytosolic tdTomato (magenta) and fixed at DIV12 to visualize TAOK1 localization and neuronal morphology. Scale bar is 20µm. Yellow inset is magnified on right to show localization of TAOK1 within dendrites and dendritic spines. Scale bar is 2µm.

**Figure S1.**
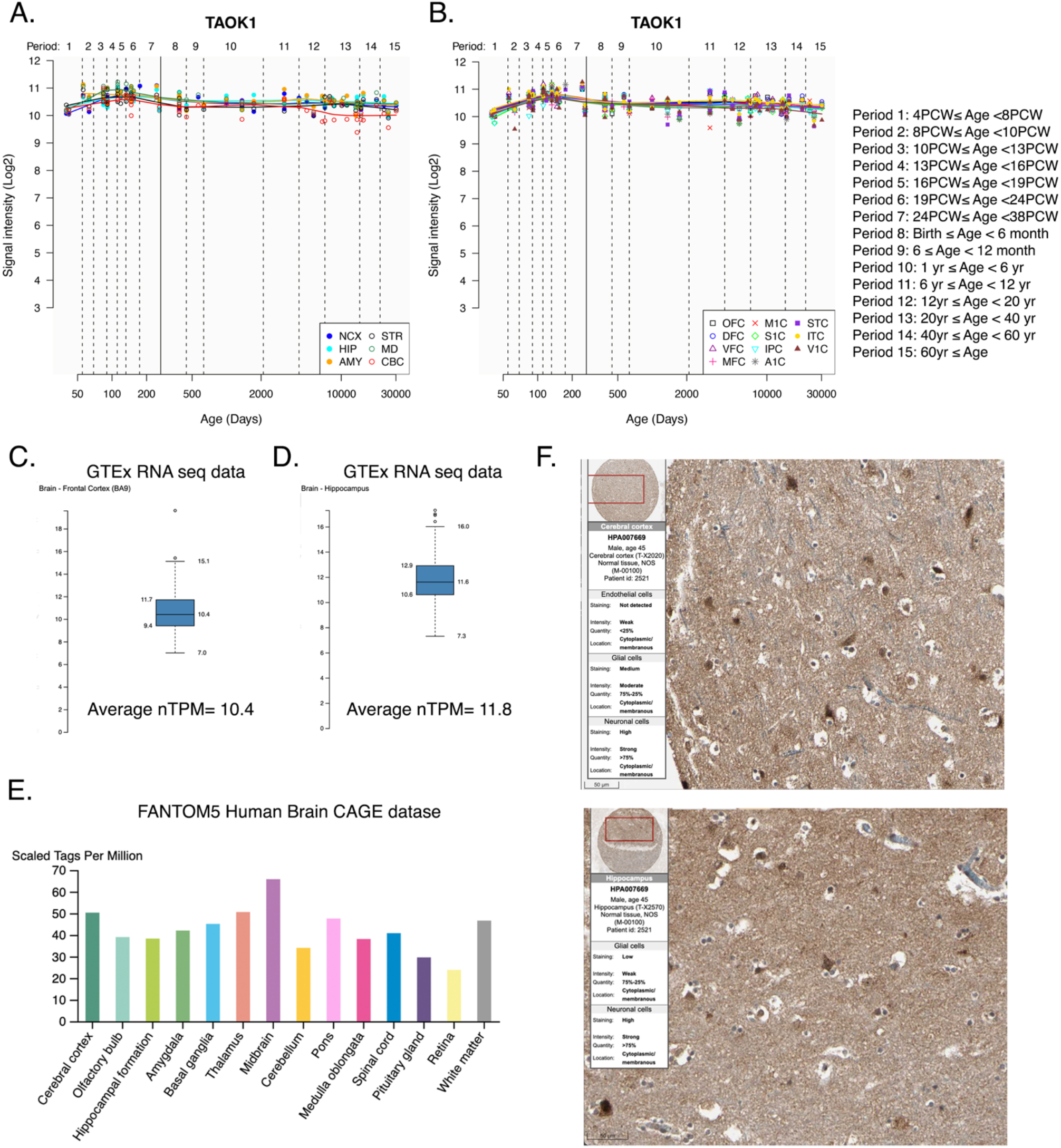
TAOK1 is expressed in human brain across development. (A) Human brain transcriptomic data for TAOK1 obtained from the public database Human Brain Transcriptome (HBT) across different stages of development. Period 1-15. Genome-wide, exon-level transcriptome data in HBT was generated using the Affymetrix GeneChip Human Exon 1.0 ST Arrays from over 1,340 tissue samples sampled from both hemispheres of postmortem human brains. Specimens range from embryonic development to adulthood (periods 1-15) and represent both males and females. Brain regions plotted are the hippocampus (HIP), neocortex (NCX), cerebellar cortex (CBC), mediodorsal nucleus of the thalamus (MD), striatum (STR), and amygdala (AMY). (B) TAOK1 expression across eleven distinct regions of the neocortex from embryonic development to adulthood is plotted. Regions include orbital prefrontal cortex (OFC), primary motor (M1) cortex (M1C), superior temporal cortex (STC), dorsolateral prefrontal cortex (DFC), primary somatosensory cortex (S1C), inferior temporal cortex (ITC), ventrolateral prefrontal cortex (VFC), posterior inferior parietal cortex (IPC), primary visual (V1) cortex (V1C), medial prefrontal cortex (MFC) and primary auditory cortex (A1C). (C-D) RNA-Seq data generated by the Genotype-Tissue Expression (GTEx) project from human tissues is reported as mean nTPM. Data from 209 samples obtained from frontal cortex BA9 (C) and hippocampal (D) region of both sexes is visualized with box plot, shown as median and 25th and 75th percentiles. Points are displayed as outliers if they are above or below 1.5 times the interquartile range. nTPM (normalized transcripts per million) values of the individual samples are presented next to the box plot. (E) Brain tissue data for TAOK1 RNA expression obtained through Cap Analysis of Gene Expression (CAGE) generated by the FANTOM5 project are reported as Scaled Tags Per Million. (F) Human Protein Atlas generated images of histological sections from normal brain tissues from cortical slices and hippocampal slices obtained by immunohistochemistry using TAOK1 antibody (HPA007669, rabbit). Antibodies are labeled with DAB (3,3’-diaminobenzidine) and the resulting brown staining indicates where an antibody has bound to its corresponding antigen. High protein expression of TAOK1 was detected in neuronal cells in both cortex and hippocampus with weak staining in glial cells. Scale bar is 50µm.

### NDD-associated mutations in TAOK1 kinase domain are catalytically dead and lead to plasma membrane protrusions

We next investigated the impact of autism and NDD-associated mutations on TAOK1. Disease associated mutations are spread out through the length of the protein encoded by *TAOK1* (Figure S2A). TAOK1 is predicted to be extremely intolerant of mutations (data from gnomAD browser), with a pLI score (Consortium et al., 2016; Zou et al., 2016) of 0.998 and the ratio of the observed/expected variants determined by O/E constraint score of 0.17. Of the 28 NDD associated mutations identified to date within TAOK1, we focused on four mutations that are clustered within the catalytic domain of TAOK1 (Figure S2A, Figure 2A). We performed in vitro kinase assays to determine the impact of these NDD mutations on the enzymatic activity of TAOK1. TAOK1 autophosphorylates itself at the S181 residue in the catalytic loop, which can be used as a measure of its kinase activity(Byeon et al., 2022). HEK293T cells were transfected with GFP-tagged wild-type or mutant TAOK1, and immunoprecipitated TAOK1 was then subjected to *in vitro* kinase assay. Probing with an antibody that recognizes the phosphorylated S181 residue, we found that wildtype TAOK1 showed robust kinase activity, however, all four NDD-associated TAOK1 mutants S111F, L167R, A219V and R269Q were catalytically dead (Figure 2B-C). Next, we expressed GFP-tagged TAOK1 constructs harboring the NDD-associated kinase-dead mutations in HEK293T cells. While wild-type (WT) TAOK1 was primarily cytosolic, the kinase-dead disease mutants were enriched at the plasma membrane (Figures 2D-E and S2B). Concomitant with their plasma membrane localization, all four NDD associated kinase-dead TAOK1 constructs led to aberrant and exuberant protrusions of the plasma membrane, while expression of the wildtype TAOK1 did not promote membrane protrusions (Figures 2D and S2B-S2C).

**Figure 2.**
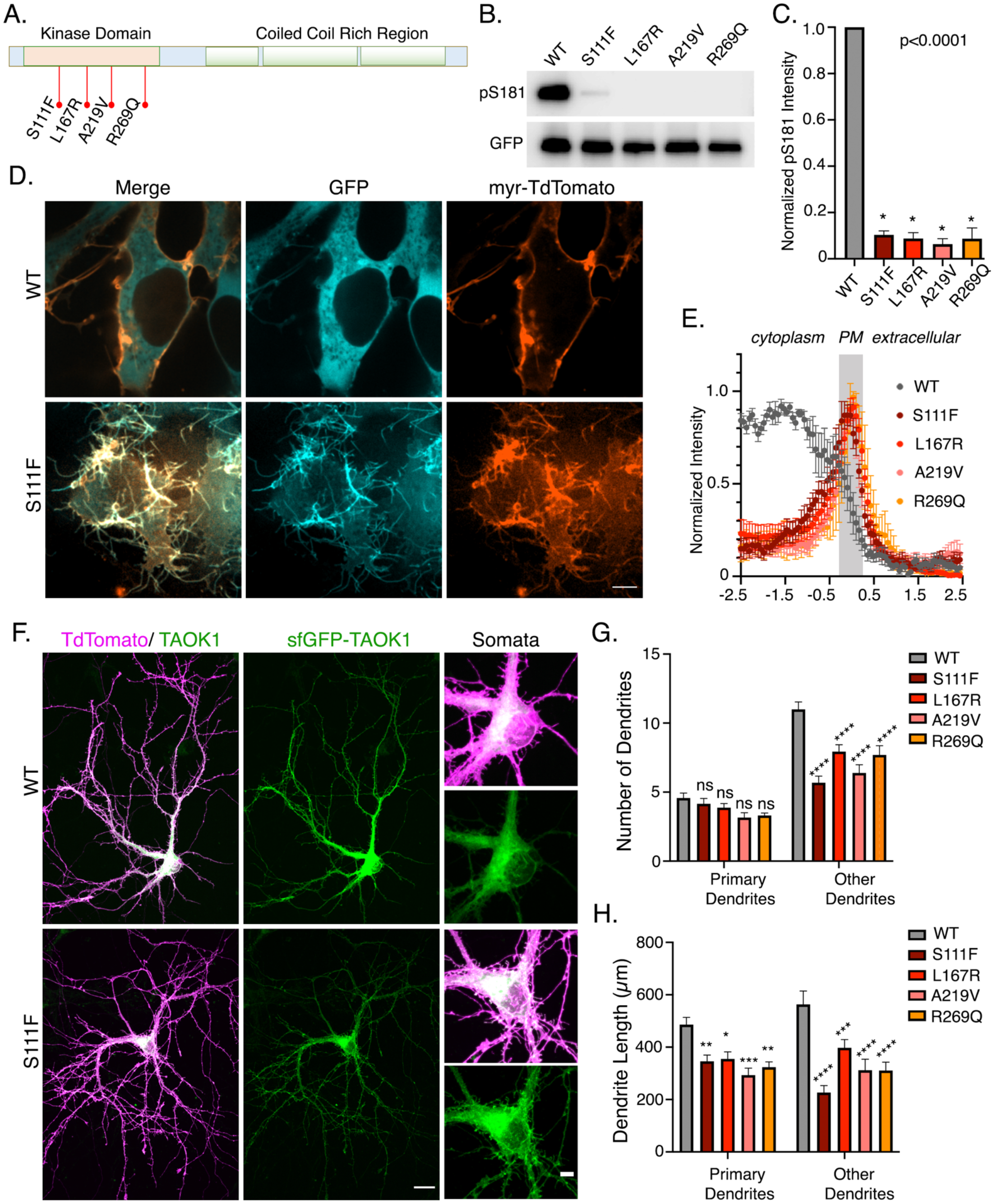
NDD associated mutations that render TAOK1 catalytically dead induce aberrant membrane protrusions and perturb normal dendrite development. (A) Schematic depicts the secondary protein structure of TAOK1, and highlights the position of the four autism-associated mutations S111F, L167R, A219V and R269Q clustered within the N-terminal kinase domain of TAOK1. (B) Western blot shows *in vitro* kinase activity of immunoprecipitated GFP-tagged wildtype (WT) TAOK1 and the four mutants as indicated. Autophosphorylation at residue S181 is used as a readout for kinase activity and amount of total TAOK1 protein is indicated by GFP blot. (C) Kinase activity as measured by intensity of pS181 signal to the GFP signal is normalized to WT TAOK1 and plotted in the bar graph. Mean values are depicted and error bars represent standard error of mean (n=3, p<0.0001). (D) Images show representative HEK293T cells expressing plasma membrane marker myr-tdTomato (red) along with either wildtype TAOK1 (WT) or NDD-associated TAOK1 mutants S111F (cyan). Single confocal plane is shown. Scale bar is 10µm. (E) Plot depicts the normalized fluorescence intensity distribution of GFP-TAOK1 wildtype and autism mutants S111F, L167R, A219V and R269Q across a 5µm line drawn across a single z plane of a confocal image such that 2.5µm is intracellular, 0µm refers to plasma membrane and the remaining 2.5µm is extracellular. Note the peak fluorescence of the mutants at the plasma membrane while the WT is mostly intracellular. Normalized mean intensity values as a function of distance are plotted and error bars represent standard error of mean (n=6 cells per condition, p<0.0001). (F) DIV11 rat hippocampal neurons expressing WT-TAOK1 or S111F ASD mutant TAOK1 (green) along with cytoplasmic td-tomato (magenta) to visualize neuronal morphology. Right column shows the magnified somatic region of the neuron to highlight the largely cytoplasmic localization of WT-TAOK1 (top) compared to the membrane associated localization of S111F mutant which leads to numerous membrane protrusions from neuronal soma. (G) Total number of primary dendrites and secondary or tertiary (others) dendrites in hippocampal neurons expression GFP-tagged WT TAOK1 or the mutants S111F, L167R, A219V and R269Q is plotted. Mean values are plotted and error bars represent standard error of mean (n>14 neurons per condition from 3 different experiments, 2-way ANOVA with multiple comparisons to mean WT value, ns= not significant and **** indicates p<0.0001). (H) Total length of primary dendrites and secondary or tertiary (others) dendrites in hippocampal neurons expression GFP-tagged WT TAOK1 or the mutants S111F, L167R, A219V and R269Q is plotted. Mean values are plotted and error bars represent standard error of mean (n>14 neurons per condition from 3 different experiments, 2-way ANOVA with multiple comparisons to mean WT value, ns= not significant, *p<0.05, **p<0.005, ***p<0.001 and **** indicates p<0.0001).

**Figure S2.**
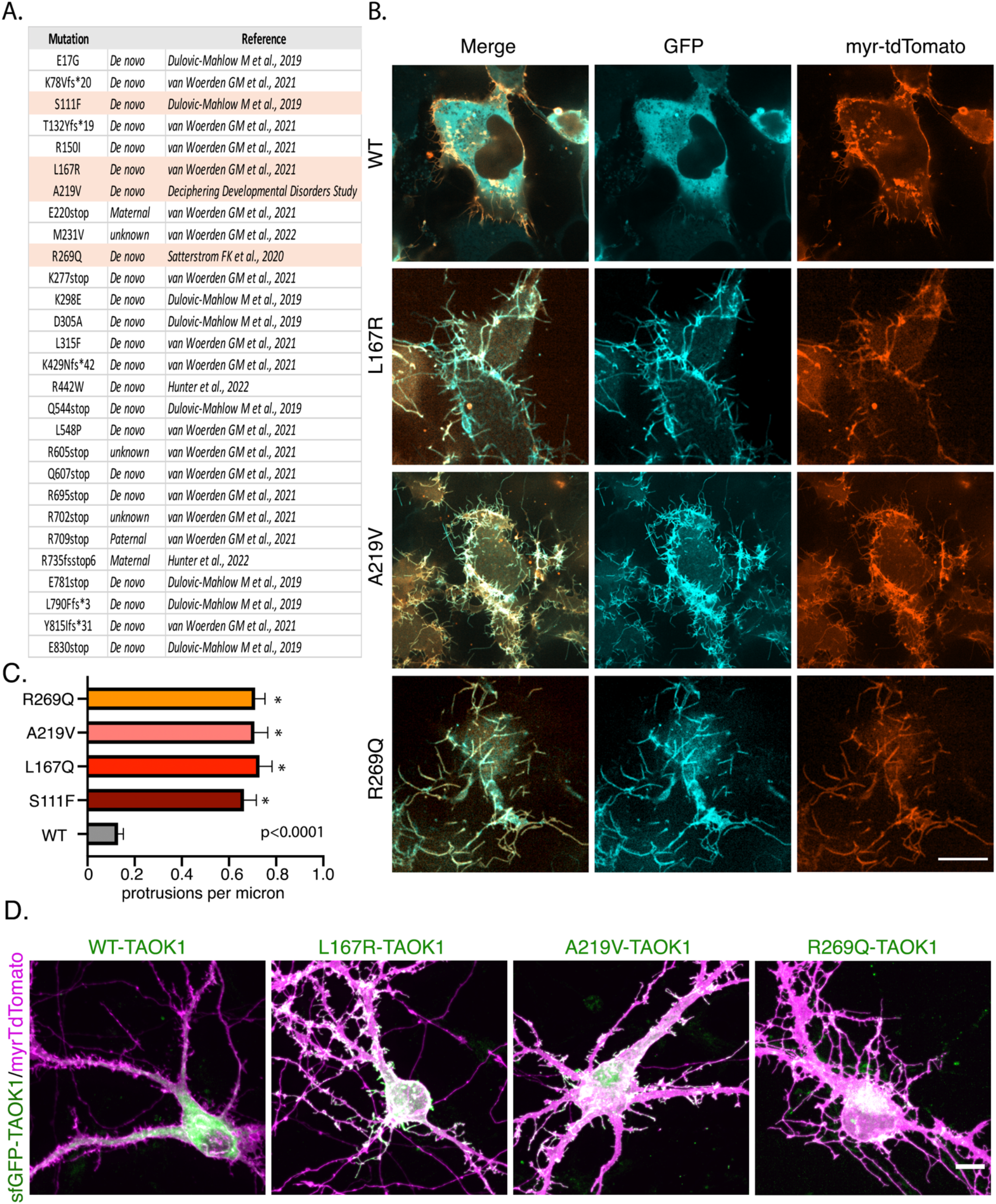
NDD associated mutations render TAOK1 catalytically dead and induce aberrant neuronal protrusions (related to main figure 2) (A) Compiled list of TAOK1 missense and frameshift mutations that have been associated with neurodevelopmental disorders. The mutations that are characterized as kinase dead are highlighted in orange. Mutations are either inherited or acquired *de novo* and have been found in both males and females. (B) HEK293T cells expressing plasma membrane marker myr-tdTomato along with either wildtype TAOK1 (WT) or autism associated TAOK1 mutants S111F, L167R, A219V and R269Q. Single confocal plane is shown. Scale bar is 10µm. (C) Bar plot depicts mean density of membrane protrusions in HEK293 cells expressing plasma membrane marker myr-tdTomato along with either wildtype TAOK1 (WT) or autism associated TAOK1 mutants S111F, L167R, A219V and R269Q. Error bars show standard error of mean, n=10 cells per condition. One way ANOVA with multiple comparisons, p<0.0001. (D) DIV11 hippocampal neurons expressing WT-TAOK1 or ASD mutants L167R, A219V and R269Q TAOK1 (green) along with cytoplasmic td-tomato (magenta) to visualize neuronal morphology. Scale bar is 10µm.

### NDD-associated mutations in TAOK1 cause defects in dendrite development

To test the functional impact of these mutations on neuronal development, we expressed wildtype TAOK1 and ASD mutant TAOK1 in hippocampal neurons. Neurons were transfected at DIV9 and then fixed at DIV11 to determine the impact of these mutations on dendritic growth. Expression of the kinase-dead NDD associated TAOK1 mutants severely perturbed neuronal development compared to WT-TAOK1 (Figure 2F-H, S2D). Similar to HEK293T cells, the kinase dead TAOK1 mutants localized to the plasma membrane and induced abnormal protrusions which were most apparent within the soma of the hippocampal neurons (Figure 2F (inset) and S2D). While the number of primary dendrites was not affected, number of secondary and tertiary dendrites were significantly reduced on expression of the four kinase-dead mutants (Figure 2G). Further, total dendrite length of both primary dendrites and secondary/tertiary dendrites was significantly decreased in neurons expressing the TAOK1 mutants as opposed to the WT TAOK1 (Figure 2H).

### TAOK1 coiled-coil helical bundle is necessary and sufficient for plasma membrane tubulation

We next probed the mechanism through which TAOK1 might localize to the plasma membrane and cause tubulations. We generated several GFP-tagged TAOK1 deletion constructs to map the minimal domain required for its plasma membrane localization (Figure 3A). When expressed in HEK293T cells, the GFP-tagged kinase domain (residues 1-320) construct was cytosolic, while the construct lacking the kinase domain (residues 321-1001) localized to the plasma membrane and led to formation of extensive membrane protrusions (Figure S3A). Further dissection revealed that the C-terminal tail (901-1001) was dispensable, and that residues 321-901 were necessary and sufficient for TAOK1 association with the plasma membrane and for generation of membrane protrusions (Figure 3B and Figure S3A-B). Cellular fractionation of HEK293T cells expressing GFP-tagged kinase domain (1-320) or coiled-coil domain (321-901), confirmed that the coiled-coil domain was primarily membrane associated while the kinase domain was cytosolic (Figure 3C-D). Live-cell imaging revealed that the membrane protrusions were stable membrane extensions that were actively extending, but rarely retracted (Figure S3B). Average length of membrane protrusions in cells expressing TAOK1 coiled-coil domain increased to 11µm while the average in control cells expressing TAOK1(1-320) was 2µm (Figure 3E). Expression of the membrane binding domain (321-901) of TAOK1 in neurons led to severe perturbation of neuronal morphology similar to that exhibited by the NDD mutants (Figure 3F), while kinase domain or C-terminal tail domains expressing constructs were localized to the cytoplasm and did not perturb neuronal morphology (Figure 3F and 3G).

**Figure 3.**
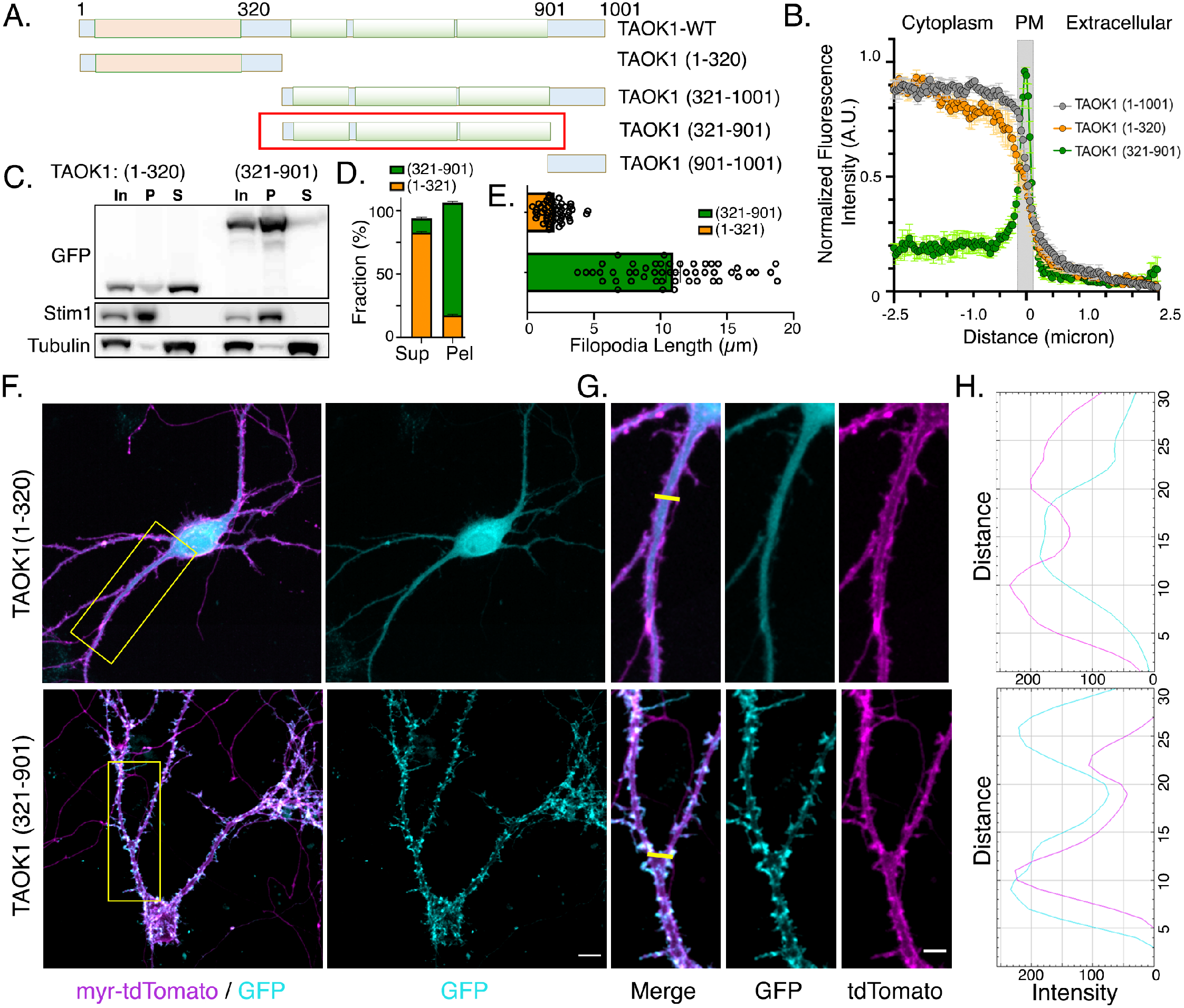
Coiled coil domain of TAOK1 binds and tubulates plasma membrane. (A) Schematic depicts the different deletion constructs generated to identify the membrane binding domain within TAOK1. The red box indicates the region necessary and sufficient for TAOK1 plasma membrane association. (B) Plot depicts the normalized fluorescence intensity distribution of full length GFP-TAOK1 (1-1001 amino acids), kinase domain (1-320 amino acids) and coiled coil domain (321-901 amino acid) across a 5µm line drawn across a single z plane of a confocal image such that 2.5µm is intracellular, 0µm refers to plasma membrane and the remaining 2.5µm is extracellular. Note the peak fluorescence of coiled coil domain (321-901) at the plasma membrane compared to the mostly intracellular intensity of WT and isolated kinase-domain. Normalized mean intensity values are plotted as a function of distance and error bars represent standard error of mean (n=10 cells per condition, p<0.0001). (C) Western blot shows the localization of GFP-tagged TAOK1 kinase domain (1-320) and coiled-coil domain (321-901) in supernatant (S) and membrane (M) fractions, respectively. Fractions were obtained by differential centrifugation at 100,000g. TAOK1 domains were detected by immunoblotting using anti-GFP antibodies. ER protein Stim1 and α-tubulin are used as controls for the membrane and supernatant fractions, respectively. (D) Percentage of GFP tagged kinase domain and coiled-coil domain present in the membrane and supernatant fractions is plotted (n=3, p<0.0001). (E) Mean length of plasma membrane protrusions in HEK293T cells expressing either the kinase domain (1-320) or the coiled coil domain (321-901) is plotted. (n=50 protrusions from 10 cells each, p<0.0001). (F) DIV11 rat hippocampal neurons expressing GFP-tagged TAOK1 kinase domain (1-320) or coiled coil domain (321-901) (cyan) along with myristoylated-tdTomato (magenta) to visualize neuronal morphology. Yellow inset marks the area magnified in F. Scale bar is 10µm. (G) Magnified dendritic region shows cytoplasmic localization of TAOK1 (1-320) and plasma membrane localization of (321-901) overlapping extensively with the membrane marker myr-TdTomato (magenta). Yellow bar marks the dendritic plane that is analyzed in G for colocalization of TAOK1 and myr-TdTomato. Scale bar is 3µm. (H) Plot profile of GFP (cyan) and myrTdTomato (magenta) fluorescence as a function of distance (in pixels) indicates high co-localization of coiled coil domain (321-901) with the plasma membrane while kinase domain (1-320) does not show colocalization.

**Figure S3.**
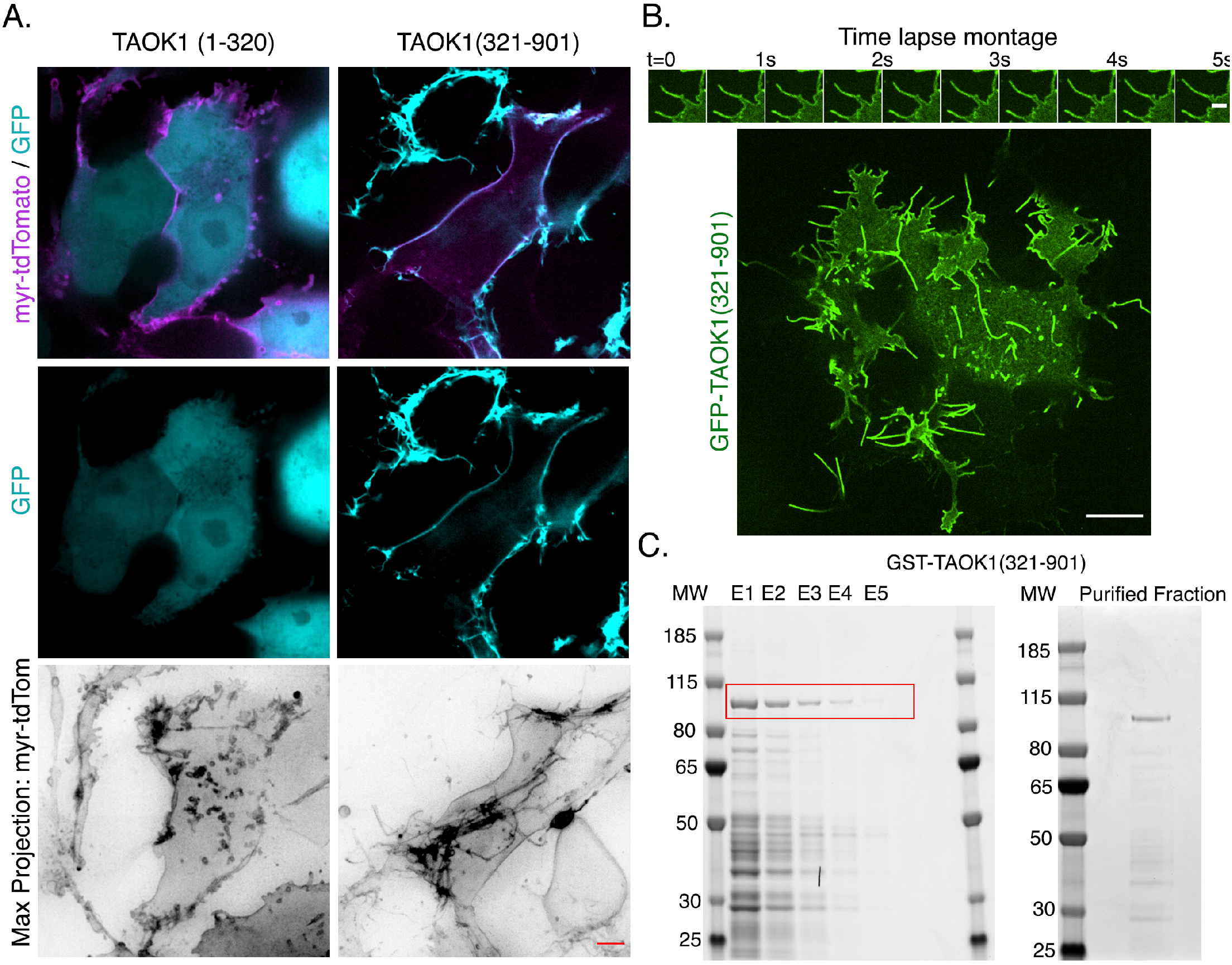
Coiled coil domain of TAOK1 binds and tubulates plasma membrane (related to Figure 3) (A) Single z-frame confocal image of HEK293T cells transfected with TAOK1 kinase domain sfGFP-TAOK1 (1-320) (left column) and helical bundle domain sfGFP-TAOK1 (right column) (cyan, bottom row) along with plasma membrane marker myristoylated-tdTomato (magenta). Maximum projected and inverted image of myr-tdTomato to visualize the membrane protrusions is shown in grayscale (bottom row). Scale bar is 5µm. (B) Montage shows membrane protrusion enriched with GFP-TAOK1 (321-901) extending in length, frames were taken every 0.5s and scale bar is 2µm. Time lapse imaging of a single confocal z-frame of HEK293T cells expressing GFP-TAOK1 (321-901), scale bar is 10µm. Yellow box shows the area of which the time-lapse montage was generated. (C) GST-TAOK1(321-901) with molecular weight 95.7 KDa was bacterially expressed and affinity purified on glutathione beads. Elution fractions E1-E5 are shown on an SDS page Coomassie stained gel (left). Fractions E2-E5 were combined and further purified through a 75KDa cut off column (right).

### TAOK1 helical bundle directly binds phosphoinositides

To gain structural insight into how the coiled-coil domain of TAOK1 might associate with plasma membrane, we used the machine learning based protein-structure prediction software, AlphaFold2.0 (Jumper et al., 2021; Tunyasuvunakool et al., 2021). The prediction revealed that the three coiled-coils fold into a triple helix bundle (gray) that makes intimate interactions with the kinase domain (green) (Figure 4A, B). We used the Adaptive Poisson-Boltzmann Solver (Jurrus et al., 2018) function of Pymol to generate an electrostatic map of surface residues of the predicted triple helix. The triple helix domain formed a gently curved crescent shaped structure with prominent separation of charged residues. The convex side of the triple helix was positively charged while the concave surface was negatively charged (Figure 4B, side view). This suggested that the triple helix could potentially directly bind negatively charged lipids in the plasma membrane through electrostatic interactions with the basic amino acids on its convex surface. To test whether the TAOK1 triple helix can directly bind lipids, we bacterially purified GST-tagged TAOK1 (321-901) protein (Figure S3C). We used the lipid overlay assay to determine whether TAOK1 triple helix could directly bind lipid molecules. We found that TAOK1 specifically bound phosphatidylinositol (4)-phosphate (PI4P), phosphatidylinositol (4,5)-bisphosphate, and phosphatidylinositol (3,4,5)-triphosphate, but did not bind phosphatidylinositol (PI) or phosphatidylethanolamine (PE) (Figure 4C-D). The PI(4,5)P2 sensor GFP-C1-PLC∂-PH (Stauffer et al., 1998) when transfected in HEK293T cells localized to the plasma membrane as expected. Consistent with the tubulating activity of TAOK1 triple helix bundle, when mCherry-TAOK1(321-901) was co-transfected with the PI(4,5)P2 sensor, we observed the development of exuberant protrusions of the plasma membrane (Figure 4E). These results collectively show that TAOK1 can directly interact with negatively charged phospholipids enriched in the plasma membrane through its triple helical domain.

**Figure 4.**
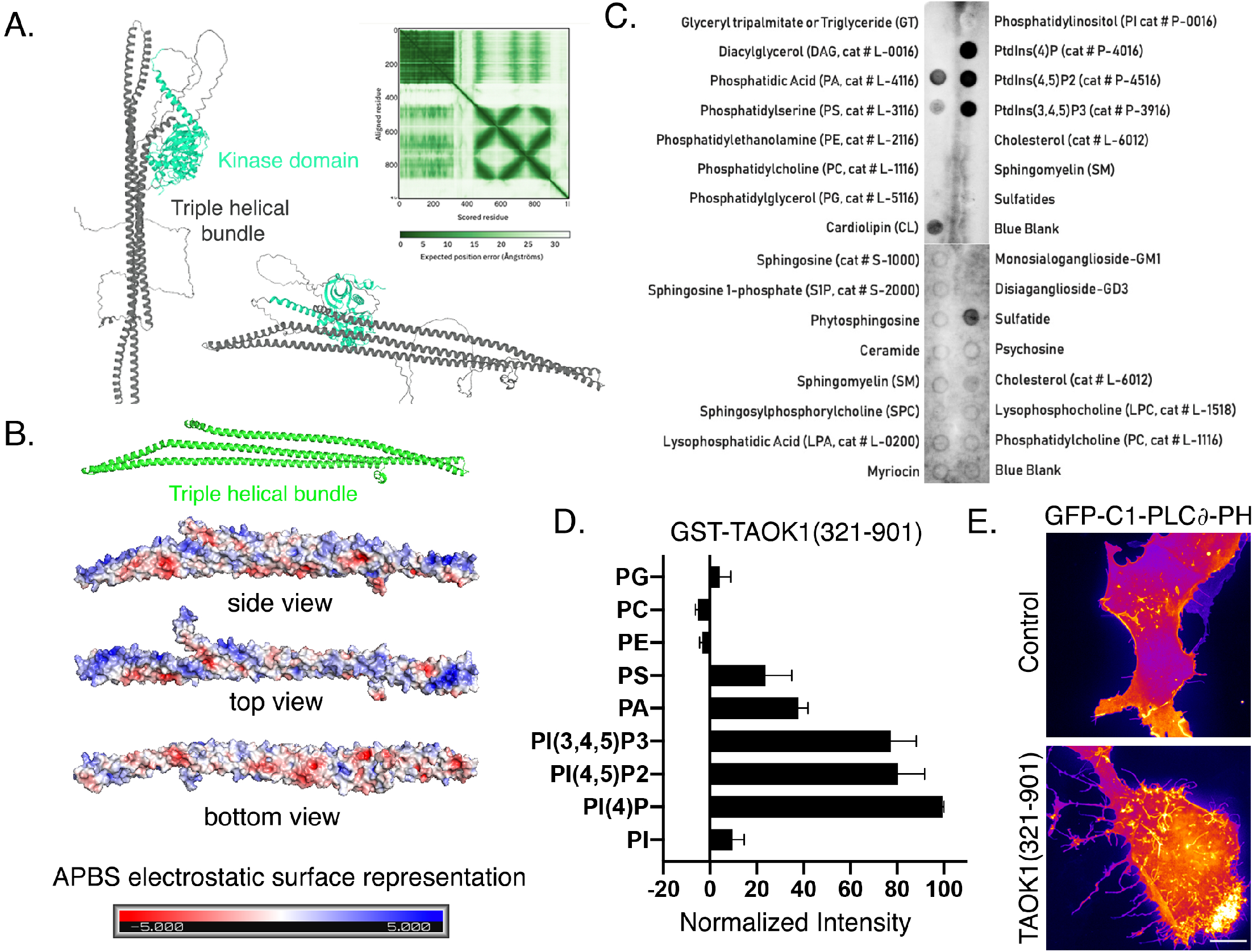
TAOK1 directly associates with phosphoinositides and sculpts the plasma membrane. (A) AlphaFold2.0 prediction of the structure of human TAOK1. The kinase domain is shown in green (residues 1-320) and the rest of protein is shown in gray (321-1001). The three coiled coil domains are predicted to fold into a triple helix. Expected position error at each residue is plotted for the entire length of the protein (top right). (B) The isolated triple helix bundle is shown in neon green. The APBS (Adaptive Poisson-Boltzmann Solver) function of Python was used on the Alphafold2.0 predicted structure of TAOK1 triple helix to create a surface electrostatic representation. The convex face of the crescent shaped protein domain (side view, top view) is highly enriched in positive charges (blue) while the concave side (side view, bottom view) is enriched in negative charged amino acids (red). The scale of charges −5 to +5 is shown as gradation in shades of red and blue. (C) Lipid overlay blots show the binding affinity of GST tagged TAOK1 triple helix (321-901) to different species of lipids dotted on the blot. Darker dots depict greater binding which was detected through antibody against GST tag. (D) Quantification of binding affinity of GST-TAOK1(321-901) with different lipids. Mean values are plotted and error bars depict stand error of mean (n=3 experiments). (E) Images shows representative HEK293T cells transfected with the PI(4,5)P2 sensor GFP-C1-PLC∂-PH along with either control (top) or mCherry-TAOK1(321-901). Co-expression of TAOK1 helical bundle leads to massive protrusions from the plasma membrane. Scale bar is 5µm.

### Catalytic activity of TAOK1 rescues plasma membrane association and tubulation

Upon discovering that TAOK1 can directly bind plasma membrane through association with negatively charged phosphoinositides, we investigated why TAOK1 kinase-dead mutants, but not the active kinase, were strongly plasma membrane-bound. One possibility was that the kinase domain autophosphorylates itself to negatively regulate membrane association. To test this hypothesis, we transfected HEK293T cells with either GFP-tagged TAOK1 (321-901) alone or coexpressed with mCherry-TAOK1(1-320) kinase domain (Figure 5A). When expressed alone TAOK1 triple helical domain localized to the plasma membrane and led to numerous protrusions, however, co-expression of the kinase domain (1-320) with the triple helical domain rescued both the membrane binding and tubulating effect of the TAOK1 triple helix (Figure 5A-B). The inactive kinase domain TAOK1 (1-320 K57A) did not rescue the membrane localization of the TAOK1 triple helix. Importantly, expression of the isolated kinase domain (1-320) rescued the membrane association of all the four kinase-dead NDD associated mutants S111F, L167R, A219V and R269Q (Figure 5C-D). Further, in presence of this active kinase-domain, TAOK1 disease mutants also failed to generate aberrant cellular protrusions (Figure 5C, D). These results strongly suggest that the TAOK1 kinase activity inhibits membrane-binding of the triple helix through the phosphorylation of key residues within the triple helical domain.

**Figure 5.**
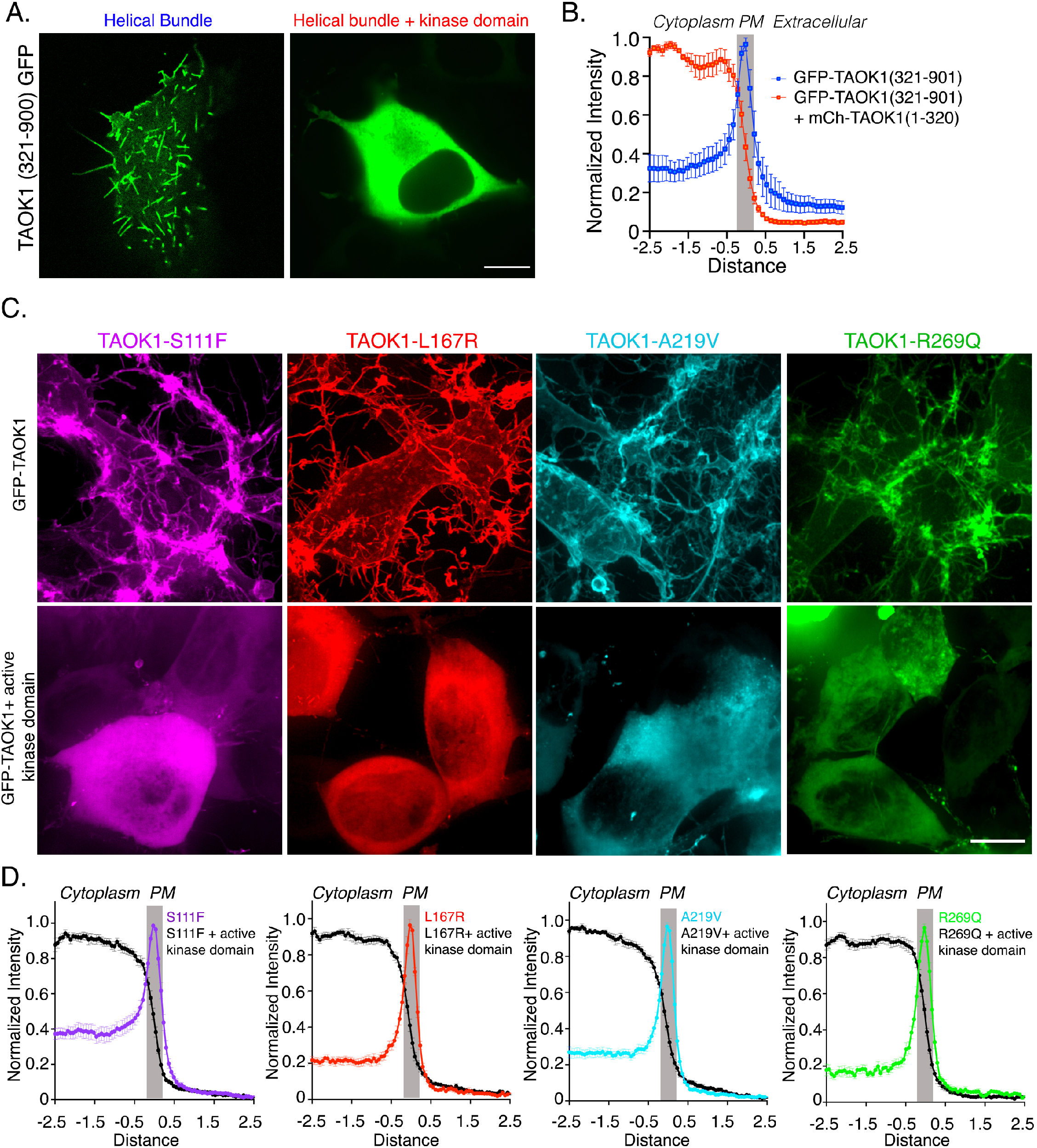
Autophosphorylation rescues aberrant plasma membrane association and tubulation by TAOK1. (A) HEK293T cells transfected with TAOK1 helical bundle domain GFP-TAOK1 (321-901) alone or with TAOK1 kinase domain mCherry-TAOK1 (1-321). Scale bar is 5µm. (B) Plot depicts the normalized fluorescence intensity distribution of GFP-TAOK1 (321-901) alone (blue) or GFP-TAOK1 (321-901) with TAOK1 kinase domain mCherry-TAOK1 (1-321) (red) across a 5µm line drawn across a single z plane of a confocal image such that 2.5µm is intracellular, 0µm refers to plasma membrane and the remaining 2.5µm is extracellular. Note the expression of kinase domain rescues the peak fluorescence of coiled coil domain (321-901) at the plasma membrane. Normalized mean intensity values as a function of distance are plotted and error bars represent standard error of mean (n=10 cells per condition, p<0.0001). (C) Representative HEK293T cells expressing NDD-associated TAOK1 mutants S111F (magenta), L167R (red), A219V (cyan) and R269Q (green) are shown in top row, and representative HEK293T cells expressing these autism mutants along with active kinase domain of TAOK1 (1-320) are shown in bottom row. Scale bar is 10µm. (D) Plot depicts the normalized fluorescence intensity distribution of TAOK1 mutants S111F (magenta), L167R (red), A219V (cyan) and R269Q (green) across a 5µm line drawn across a single z plane of a confocal image such that 2.5µm is intracellular, 0µm refers to plasma membrane (gray box) and the remaining 2.5µm is extracellular. Note the expression of kinase domain rescues the peak fluorescence of autism mutants at the plasma membrane (all plots, black curve). Normalized mean intensity values as a function of distance are plotted and error bars represent standard error of mean (n=6 cells each per condition, p<0.0001).

### Proteomic identification of residues that control TAOK1 membrane association

We hypothesized that TAOK1 induces conformational changes in its helical membrane-binding domain through autophosphorylation, thereby negatively regulating membrane binding and tubulation. To test this, we first used mass spectrometry to identify residues that TAOK1 might phosphorylate in order to regulate its membrane association. GFP-tagged wildtype TAOK1 and kinase-dead TAOK1 (L167R) were expressed in HEK293T cells and immunoprecipitated using anti-GFP antibodies. Following extensive salt washes to remove co-immunoprecipitating proteins, immunoprecipitated TAOK1 was subjected to an in vitro kinase assay to allow for autophosphorylation. Several phosphorylated residues were identified by mass spectrometry for WT kinase (Figure 6A-B). TAOK1/2 are known to prefer phosphorylation at threonine residues (Miller et al., 2019; Yadav et al., 2017), hence we focused on the three threonine phosphorylation sites T440, T443 and T747 identified within the helical bundle (Figure 6B). To test which sites were important for regulation of TAOK1 membrane-binding, we generated phosphomutant constructs that abolished phosphorylation at the identified sites by mutating the threonine to an alanine residue. Mutating sites T440A/T443A led the kinase-active TAOK1 to now bind the plasma membrane and generate extensive membrane tubules, while no apparent effect was observed for T747A mutation (Figure 6C and D). These data provide direct evidence that autophosphorylation by TAOK1 at the T440 and T443 residues is critical for regulating membrane association and outward protrusions. To test whether one or both residues were required to modulate TAOK1 membrane-association, we generated a GFP-tagged construct 441-900, which lacks the phosphorylation site T440. GFP-TAOK1(441-900) when expressed in HEK293T cells localized to the plasma membrane and generated outward membrane protrusions (Figure 6E-F). While expression of the active kinase domain mCherry-TAOK1(1-320) along with GFP-TAOK1(321-900) inhibited its plasma membrane association, expression of the kinase domain (1-320) failed to rescue the membrane localization and tubulating properties of GFP-TAOK1(441-900). These data demonstrate that the triple helix lacking the T440 residue loses its ability to dissociate from the plasma membrane by TAOK1 autophosphorylation. Further, these results provide direct evidence for TAOK1 mediated mechanisms for remodeling the plasma membrane in an activity dependent manner (Figure 6G).

**Figure 6.**
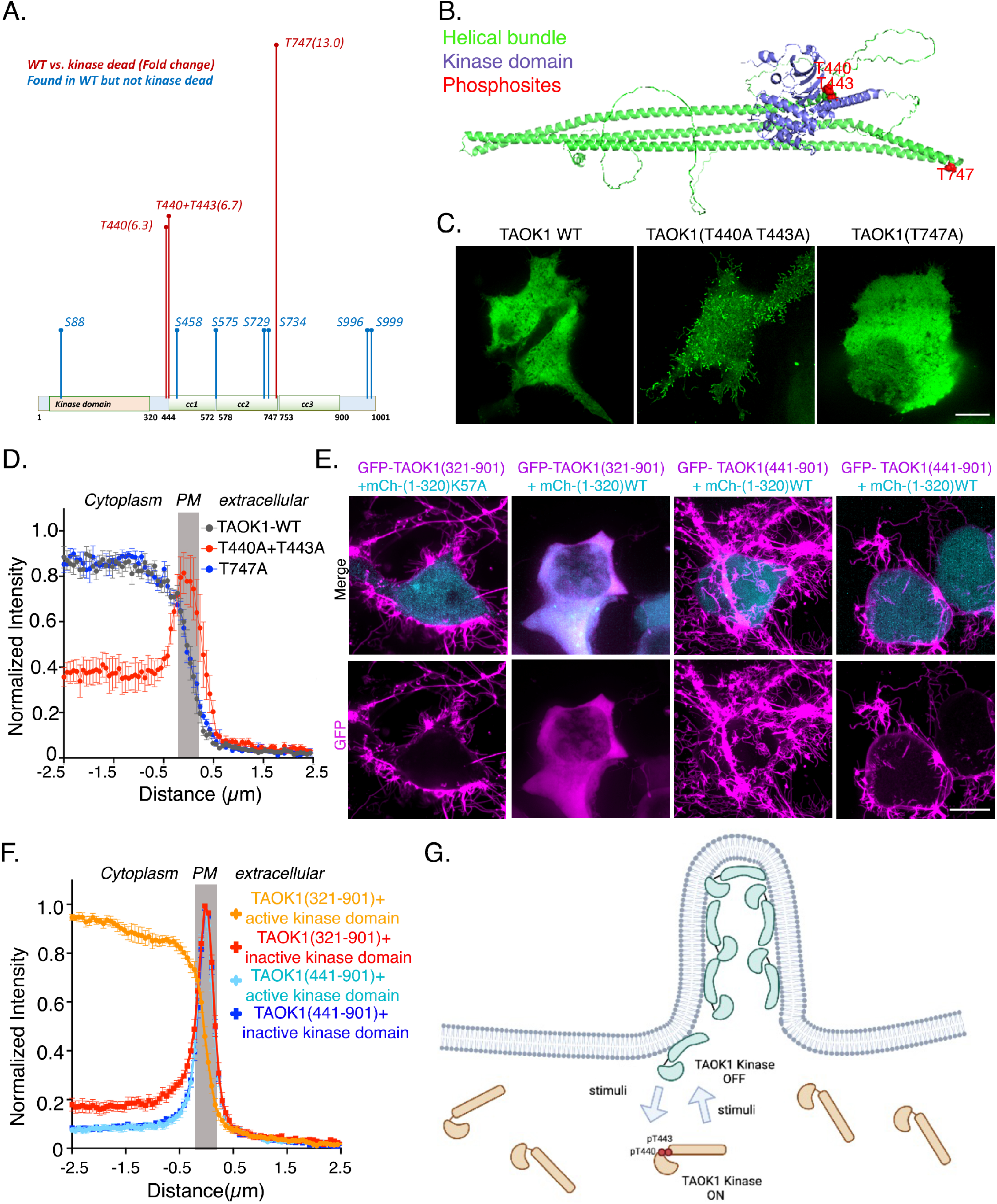
Phosphorylation at residues T440/T443 provide mechanism for autoregulation of TAOK1 membrane association and remodeling function. (A) Schematic shows the phosphorylation sites identified by mass spectrometry using immunoprecipitated WT-TAOK1 and kinase dead (L167R) TAOK1 was used as negative control. The fold change increase in WT over kinase-dead control for the three phosphothreonine sites (red) identified is shown in parenthesis after the phosphosite. T440 (6.3 fold higher in WT), T443 (6.7 fold higher in WT) and T747 (13 fold higher in WT). Seven serine phosphorylation sites shown in blue were only identified in WT and not kinase dead control. Localization of the identified phosphosites is shown by overlay with the secondary protein structure of TAOK1. (B) Position of the phospho-Threonine sites is indicated in the 3D predicted structure of TAOK1. Both T440 and T443 are in close proximity to where the kinase domain contacts the triple helical bundle. The T747 site is localized at the loop where 2 coil coils bend. Kinase domain is shown in blue, helical bundle in green and phospho-sites in red. (C) Representative confocal images of HEK293T cells transfected with TAOK1-WT (left), double phosphomutants TAOK1-(T440A+T443A), and phosphomutant TAOK1-(T474A) (right). Scale bar is 10µm. (D) Plot depicts the normalized fluorescence intensity distribution of TAOK1-WT (gray), double phosphomutants TAOK1-(T440A+T443A) (red), and phosphomutant TAOK1-(T474A) (blue) across a 5µm line drawn across a single z plane of a confocal image such that 2.5µm is intracellular, 0µm refers to plasma membrane (gray box) and the remaining 2.5µm is extracellular. Note the expression of double mutant T440A+T443A renders the kinase active full-length TAOK1 trapped on the plasma membrane as indicated by the peak fluorescence at the plasma membrane. Normalized mean intensity values as a function of distance are plotted and error bars represent standard error of mean (n=6 cells each per condition, p<0.0001). (E) Representative confocal images of HEK293T cells transfected with GFP-tagged TAOK1-(321-901) and GFP-tagged TAOK1-(441-901) in magenta, along with either the active isolated kinase domain mCh-TAOK1-(1-320WT) or the inactive kinase domain mCh-TAOK1-(1-320 K57A) shown in cyan. Scale bar is 10µm. (F) Plot depicts the normalized fluorescence intensity distribution of GFP-tagged TAOK1-(321-901) and TAOK1-(441-901) expressed along with either the active isolated kinase domain mCh-TAOK1-(1-320WT) or the inactive kinase domain mCh-TAOK1-(1-320 K57A) across a 5µm line drawn across a single z plane of a confocal image such that 2.5µm is intracellular, 0µm refers to plasma membrane (gray box) and the remaining 2.5µm is extracellular. Note the expression of kinase domain rescues the TAOK1-(321-901) trapped on the plasma membrane (orange curve) but fails to do so for the TAOK1-(441-901) as indicated by the peak fluorescence at the plasma membrane (cyan curve). Normalized mean intensity values as a function of distance are plotted and error bars represent standard error of mean (n=6 cells each per condition, p<0.0001). (G) Working model of TAOK1 kinase as a membrane sculpting kinase autoregulated by its kinase activity through phosphorylation at critical residues T440 and T443. TAOK1 shuttles between its active/cytosolic state (orange) and inactive/membrane bound state (green), which is regulated by phosphorylation. NDD associated mutations (S111F, L167R, A219V and R269Q) render TAOK1 catalytically dead and trap them permanently in their membrane bound state where it induces aberrant membrane extensions and dendritic defects, thus contributing to pathology in TAOK1 associated neurodevelopmental disorder.

## DISCUSSION

Membrane associating proteins that can sense and/or induce membrane curvature provide a direct mechanism for sculpting neuronal morphology (Carlson et al., 2011; Chatzi and Westbrook, 2021; Coutinho-Budd et al., 2012; Dharmalingam et al., 2009; Guerrier et al., 2009). Aberrations or dysregulation of plasma membrane remodeling during neuronal differentiation, migration or synaptogenesis can underlie the pathogenesis of neurodevelopmental disorders. In this study, we report that the autism and NDD-associated gene *TAOK1* encodes a plasma membrane sculpting kinase. We show that TAOK1 associates with membrane through direct interaction of its coiled-coil domain with phosphoinositides enriched in the plasma membranes. We provide evidence that this ability to associate with membrane leads to emergence of outward protrusions of the plasma membrane. TAOK1 autophosphorylation controls its plasma membrane binding and tubulation properties. Dysfunction in TAOK1 catalytic activity leads to failure in its disengagement from the plasma membrane causing aberrant neuronal membrane protrusions that perturb neuronal morphology. We characterized four distinct NDD mutations that were clustered in the kinase domain, and show that all four mutations render TAOK1 kinase-dead and result in uncontrolled membrane tubulation. In hippocampal neurons, expression of these TAOK1 mutants cause severe defects in dendritic growth. These findings provide a direct mechanism through which NDD-associated mutations in TAOK1 perturb neuronal development.

While there is only one TAO gene in invertebrates, vertebrates express three TAO genes, TAOK1, TAOK2 and TAOK3. Even though the kinase domain is highly conserved, these three TAO kinases have divergent C-terminal tails that confer them with specific and distinct functions. Both TAOK1 and TAOK2 are associated with neurodevelopmental disorders (Nourbakhsh and Yadav, 2021). Recently, we identified TAOK2 as a transmembrane kinase that tethers the endoplasmic reticulum to microtubules (Nourbakhsh et al., 2021). This study provides the first evidence that TAOK1 binds phosphoinositides in the plasma membrane and generates membrane protrusions. To our knowledge, this function makes TAOK1 unique, not only among the TAO family but among all protein kinases. It is notable that among the three mammalian TAO kinases, TAOK1 is the most evolutionarily similar to the fly dTao and *C. elegans* kin-18, therefore, it is likely that the fly and worm Tao kinase are also plasma membrane sculpting proteins, a possibility that needs future experimental investigation. For example, loss of Tao in flies induces ectopic synapses resulting in miswired connections that alter behavioral responses (Tenedini et al., 2019). This fine-tuning of synaptic connection by Tao kinase during neuronal network development might depend on its ability to remodel the plasma membrane. The Taok1 ortholog in *C. elegans*, kin-18 (Berman et al., 2001), controls contractility and polarity of the early embryo. Kin-18, interestingly, localizes to the cortex and controls Rho localization to the membrane (Spiga et al., 2013). In light of our results, it is conceivable that kin-18 directly binds the plasma membrane in embryos and regulates polarity and early embryo contraction in *C. elegans*.

We show that the catalytic activity of TAOK1 acts as a functional switch. While catalytically active TAOK1 is cytoplasmic, in its ‘kinase-off’ state TAOK1 binds to and tubulates the plasma membrane. Our data suggest that the endogenous TAOK1 exists in both its “kinase on-cytoplasmic” and “kinase off-membrane bound” states. While our study has identified key residues in TAOK1, that are autophosphorylated to control this on-off switch, further studies are needed to understand TAOK1 autoregulation and the physiological stimuli during neuronal development that mediate phosphorylation of these residues. Our study has found that ASD mutations in TAOK1 protein that render the kinase catalytically dead cause TAOK1 to be trapped in its “kinase off-membrane bound” state thereby generating aberrant plasma membrane protrusions. We show that the disease associated aberrant protrusions can be rescued by restoring the kinase activity of TAOK1, underscoring the potential for design of small molecule allosteric kinase activators to rescue the kinase from its membrane trapped state.

From a structural perspective, TAOK1 provides important future avenues to understand how its membrane binding is regulated. Using the Alphafold2.0 predicted structure, we show that surface residues lining the convex side of the triple helix are positively charged. These likely mediate electrostatic interactions with the negatively charged phospholipids in the plasma membrane. Based on the structural prediction, residues T440 and T443 lie in the loop connecting the triple helix to the kinase domain where it is plausible for an intramolecular phosphorylation to occur. Several potential mechanisms for regulating membrane binding could be T440/T443 phosphorylation induced conformational switch, dimerization or oligomerization as seen in BAR domain proteins, or steric hinderance by the kinase domain in its active versus inactive states. Comparative structural analyses of TAOK1 active and inactive states are needed to obtain further molecular insights into its autoregulation. Our data suggests that TAOK1 can regulate its membrane sculpting function through autophosphorylation, a property that is quite unique to TAOK1 compared to other membrane tubulating proteins such as I-BAR proteins. I-BAR domains from different proteins utilize partially distinct mechanisms to deform the plasma membrane and generate filopodia, and can also create signaling platforms at synapses (Saarikangas et al., 2009). Whether the plasma membrane sculpting triple-helix bundle of TAOK1 is structurally and functionally similar to the inverse BAR domain proteins, or if it represents an entirely new class of membrane remodeling proteins remains to be tested. Our results reveal the autism gene TAOK1 as the first membrane sculpting kinase yet identified, and provides the neurobiological basis for how TAOK1 dysfunction contributes to neurodevelopmental disorders.

## MATERIAL AND METHODS

### Cell culture and maintenance

HEK293T were grown in DMEM media (Thermo Fisher, Gibco) with 10% fetal bovine serum (Axenia BioLogix) and 1% Pen-Strep (Invitrogen). Cells were maintained at 5% CO2 and 37°C and passaged every 3-4 days. Primary neurons were obtained from dissociating hippocampi obtained from E18 Sprague-Dawley rat (Envigo-Harlan Lab) embryos, trypsin-dissociated, and plated at a density of 150,000 neurons per 18 mm glass coverslips (Fisher) or 450,000 neurons per 35 mm glass-bottom dish (MaTek), which were coated with 0.06 mg/mL poly-D-lysine (Sigma) and 0.0025 mg/mL laminin (Sigma). Plating media containing heat-inactivated 10% fetal bovine serum (HyClone), 0.45% dextrose, 0.11 mg/mL sodium pyruvate, 2 mM glutamine in MEM with Earle’s BBS was used. Media was changed 4hr after seeding neurons into maintenance media consisting of Neurobasal Media (Invitrogen), 2% B27 (Invitrogen), 100 units/mL penicillin and 100 mcg/mL streptomycin and 0.5 mM glutamine. Half of the media was replaced with new maintenance media every 3–4 days. For exogenous gene expression, neurons were transfected with 0.5–1.0µg plasmid DNA per coverslip in a 12-well plate (or 2–3µg DNA/35 mm dish) using Lipofectamine-2000 (Invitrogen) following manufacturer’s guide.

### Molecular cloning

Full length human TAOK1 was obtained from Transomics Technologies in PCR-XL-Topo plasmid and inserted into sfGFP-C1 (Plasmid #54579, Addgene), using the EcoRI and MfeI sites. The resulting N-terminally sfGFP-tagged TAOK1 construct (sfGFP-TAOK1) was fully sequenced (Genewiz, Azenta Life Sciences) and used as template to generate the ASD associated mutants (Genewiz). All constructs were confirmed by sequencing for accuracy. Domain deletion mutants were subcloned from sfGFP-TAOK1 using restriction enzymes EcoRI and MfeI (New England Biolabs) and assembled using NEBuilder HiFi DNA Assembly-New England BioLabs. Resultant plasmids were verified by sequencing. GST-TAOK1(321-901) was generated by subcloning from the sfGFP-TAOK1 into the pGEX4T1 vector using restriction enzymes NotI and SalI.

### Confocal microscopy

All live and fixed cell imaging was performed on a Nikon Ti2 Eclipse-CSU-X1 confocal spinning disk microscope equipped with four laser lines 405nm, 488nm, 561nm and 670 and the Nikon Elements software. A sCMOS Andor camera was used for image acquisition. The microscope was caged within the OkoLab environmental control setup enabling temperature and CO2 control during live imaging. Imaging was performed using Nikon 1.49 100x, Apo 60X, or 40X oil objectives. All image analyses were done using the open access Fiji software.

### Immunofluorescence and western blotting

All cells including neurons were fixed using warm 4% paraformaldehyde with 4%sucrose for 20 min at room temperature, followed by 3 washes with phosphate-buffered saline (PBS). One-hour incubation with blocking buffer (4% Normal donkey serum, 0.2M Glycine (pH 7.4) and 0.1% TritonX-100, in PBS) was followed by overnight incubation with primary antibody at 1:1000 dilution in blocking buffer at 4°C. After four 5min washes in PBS, cells were incubated with secondary antibody at 1:1,000 dilution in blocking buffer for 3hr at room temperatur or overnight at 4°C. Coverslips were washed and then mounted onto microscope glass slides with Fluoromont-G™(EMS). Samples for western blot analysis were treated with NuPAGE™ LDS Sample Buffer (4X) (Invitrogen) with 2-Mercaptoethanol at 5% and subsequently heated for 10 min at 95°C. Samples were electrophoresed on NuPAGE 4–12% Bis-Tris Polyacrylamide gels (Invitrogen) with NuPAGE MOPS running buffer (Invitrogen). Western blots were transferred to PVDF Membrane Immobilon®-P (Millipore-Sigma) with Transfer buffer (25mM Tris, 192mM Glycine, 20% (v/v) Methanol). Transferred blots were blocked in 5% non-fat dry milk and subjected to primary antibody and HRP conjugated secondary antibody before visualization with Pierce SuperSignal™ West Pico PLUS Chemiluminescent Substrate (Thermo Fisher). Western blot images were obtained using the ChemiDoc Imaging System (Bio-Rad Laboratories).

### Protein purification

TAOK1 C-terminal amino acids 321-901 were cloned into pGEX4T1 vector and transformed into BL21 E. coli (NEB) to express N-terminally GST-tagged TAOK1(321-901). A 25ml starter culture grown from a single colony overnight was used to inoculate 1L LB media, and grown to an O.D, of 0.6 at 37°C. Protein expression was induced by IPTG at a final concentration of 0.4mM for 12h at 18C. Bacteria were collected by centrifugation for 15min at 6000g, washed with ice cold PBS, and the pellet was resuspended in ice cold lysis buffer (50mM Tris pH 8.0, 5mM EDTA, 150mM NaCl, 20% glycerol, 5mM DTT, Protease Inhibitors (Roche, cOmplete tabTM) and PMSF). Addition of 4mg lysozyme (Sigma) was followed by 30min incubation with 0.5% TritonX-100 and sonication on ice. Supernatant was collected by centrifugation for 60min at 25,000g, and then incubated with Pierce™ GST Agarose for 1 h. Beads were washed with wash buffer (PBS + 1mM DTT + 0.1% Tween20) followed by wash buffer without detergent. Bound protein was eluted and collected in fractions by glutathione elution buffer at pH8.0 (50mM Tris pH8.0, 250mM KCl, 1mM DTT, 10% glycerol and 30mM glutathione).

### Immunoprecipitation kinase assay

HEK293T cells transfected with N-terminally sfGFP-tagged indicated variants of TAOK1 were lysed with HKT buffer (25mM HEPES pH7.2, 150mM KCl, 1% Triton X-100, 1mM DTT, 1 mM EDTA, Protease Inhibitors (Roche, cOmplete). Lysate was precleared with Pierce™ Protein G Agarose and immunoprecipitated with anti-GFP antibodies (Roche Cat. No. 11814460001) bound on protein G agarose. Beads were washed once with HKT, once for 10 min with HKT with 1M NaCl, and washed once with HK buffer (25mM HEPES pH 7.2, 150mM KCl, 1mM DTT, 1 mM EDTA, Protease Inhibitors (Roche, EDTA-free cOmplete tabTM)). Beads were washed twice with the Kinase Buffer (25mM Tris pH 7.5, 10mM MgCl2, 1mM DTT) prior to the in vitro kinase assay. Kinase assay was then performed by incubating with 0.5mM ATP and Halt™ Protease and Phosphatase Inhibitor Cocktail (Thermo Fisher) for 60min at 30°C on a shaking heat block. Samples were then subjected to western blot analysis detailed above to detect TAOK2 autophosphorylation by rabbit antibody against pS181 (R&D Systems, PPS037).

### Lipid overlay assay

Nitrocellulose-immobilized phospholipids (assay strips Cat. p6002; Echelon Biosciences) were blocked by incubation for 60min with 3% fatty-acid free BSA in TBST (137mM NaCl, 2.37mM KCl, 19mM Tris base, 0.1% Tween 20). Incubations were carried out at room temperature. Fresh blocking buffer containing 0.5μg/ml GST-TAOK1(321-901) purified protein was added for 60min incubation with the lipid strips. Lipid strips were washed five times with TBST, and then incubated for 1hr with a 1:1000 dilution of GST-Tag antibody (MA4-004, Thermo Fisher) in blocking buffer. The membrane was washed 5 times and incubated with a 1:10,000 dilution of horseradish peroxidase-conjugated anti-mouse antibodies in blocking buffer. Bound antibodies were detected by chemiluminescence with the SuperSignal West Pico Chemiluminescent Substrate (ThermoScientific, #34080) and imaged on the ChemiDoc Imager (BioRad).

### Proteomics and mass spectrometry

HEK293T cells in confluent 6-well plates were transfected with WT or kinase dead TAOK1 GFP-tagged constructs. After 24hr, cells were lysed in HKT buffer, immunoprecipitated using 50µl packed Protein G beads bound with anti-GFP antibodies (Roche), and in vitro kinase assay was performed on these samples as described above. To denature the proteins, urea was added to the samples to bring it to final 6M urea concentration followed by addition of TCEP (Tris(2-carboxyethyl) phosphine) to 1mM final concentration. Samples were incubated at room temperature for 30min on a slow thermomixer. CAM (chloroacetamide) was added to a final concentration of 3mM and incubated for 10min with slow mixing. Following alkylation, excess CAM was quenched by adding TCEP and pH was adjusted to 8.0. Denaturing was followed by on bead proteolytic digestion using LysC (MS grade, Wako JP)) for 2hr at 37°C on a thermomixer. Reaction mixture was diluted 3 times to bring down the Urea concentration to <2M with TEAB (Triethylammonium bicarbonate). Further digestion by Trypsin (MS grade, Pierce) was performed overnight at 37°C on a thermomixer. Digestion was stopped by addition of TFA (to final 1%). Peptide samples were then desalted on C18 StageTips and eluted for Mass spectrometric identification. Peptide samples were separated on an Thermo Easy-nLC1200 (Thermo Fisher Scientific) using 20 cm long fused silica capillary columns (100 μm ID, laser pulled in-house with Sutter P-2000, Novato CA) packed with 3μm 120 Å reversed phase C18 beads (Dr. Maisch, Ammerbuch, DE). Liquid chromatography (LC) solvent A was 0.1% (v/v) aq. acetic acid and LC solvent B was 20% 0.1% (v/v) acetic acid, 80% acetonitrile. The LC gradient was 100 min long with 5−35% B at 300 nL/min. MS data was collected with a Thermo Fisher Fusion Lumos Tribrid mass spectrometer. Data-dependent acquisition with a R=60,000 Orbitrap MS1 and R=30,000 Orbitrap MS2 scans collected with a max inject time of 100 msec and a duty cycle of 2 sec. Data .raw files were analyzed by MaxQuant/Andromeda (Cox et al., 2011) version 1.6.2.6 using protein, peptide and site FDRs of 0.01 and a score minimum of 40 for modified peptides, 0 for unmodified peptides; delta score minimum of 17 for modified peptides, 0 for unmodified peptides. MS/MS spectra were searched against the UniProt human database (updated Jan 2021). MaxQuant search parameters: Variable modifications included Oxidation (M) and Phospho (S/T/Y). Carbamidomethyl (C) was a fixed modification. Max. missed cleavages was 2, enzyme was Trypsin/P and max. charge was 7. The initial search tolerance for FTMS scans was 20 ppm. Label-free quantification (LFQ) of MS intensities from five process replicates/condition were processed with Perseus. Significance was calculated by a two-sample Student’s t-test.

### Statistics

Statistics were performed in GraphPad software Prism9.0 with the exception of mass spectrometry data for which MaxQuant Perseus software was used. Multiple groups were analyzed using ANOVA, and two group comparisons were made using unpaired t-test unless otherwise stated. A p-value less than 0.05 was deemed significant. All experiments were done in triplicate, and experimental sample size and p values are indicated with the corresponding figures.

## ACKNOWLEDGEMENT

We are grateful to funding from National Institute of Mental Health R01MH121674 to SY Mass spectrometry was supported by instrumentation and funding to SEO by R01GM129090. This work used an EASY-nLC1200 UHPLC and Thermo Scientific Orbitrap Fusion Lumos Tribrid mass spectrometer purchased with funding from a National Institutes of Health SIG grant S10OD021502 (S-E.O.). The content is solely the responsibility of the authors and does not necessarily represent the official views of the National Institutes of Health.

## AUTHOR CONTRIBUTIONS

All experiments were performed by NB unless otherwise stated. NB and SY analyzed the data. TS performed mass spectrometry experiments and did the MS data analyses. SEO supervised the mass spectrometry and MS data analyses. SY designed and supervised all experiments, obtained funding and wrote the manuscript.

